# Coherence-based spectro-spatial fillters for stimulus features prediction from electrocorticographic recordings

**DOI:** 10.1101/481572

**Authors:** Jaime Delgado, Andy Christen, Stephanie Martin, Brian N. Pasley, Robert T. Knight, Anne-Lise Giraud

## Abstract

The traditional approach in neuroscience relies on encoding models where brain responses to different stimuli are related to the latter to establish reproducible dependencies. To reduce neuronal and experimental noise, brain signals are usually averaged across trials to detect reliable and coherent brain activity. However, neural representations of stimulus features can be spread over time, frequency, and space, motivating the use of alternative methods that relate stimulus features to brain responses. We propose a Coherence-based spectro-spatial filter method that reconstructs stimulus features from intracortical brain signals. The proposed method models trials of an experiment as realizations of a random process and extracts patterns that are common across brain signals and the presented stimuli. These patterns, originating from different recording sites, are then combined (spatial filtering) to form a final prediction. Our results from three different cognitive tasks (motor movements, speech perception and speech production), concur to show that the proposed method significantly improves the ability to predict stimulus features over traditional methods such as multilinear regression with distributed lags and artificial neural networks. Furthermore, analyses of the model parameters show anatomical discriminability for execution of different motor movements. This anatomical discriminability is also observed in the perception and production of different words. These features could be exploited in the design of neuroprosthesis, as well as for exploring normal brain functioning.

## 1. Introduction

The traditional approach to investigate brain functions involves the presentation of different stimulus and the analysis of evoked brain response properties. The latter are collected either through non-invasive (e.g., electroencephalography [EEG], magnetoencephalography [MEG] or functional magnetic resonance imaging [FMRI]) or invasive (e.g., electrocorticography [ECoG]) recording techniques. Independent of the method used, the measured signals contain background noise arising from other biological processes and environmental interferences that need to filtered out or attenuated. Several preprocessing methods such as signal filtering or referencing can serve to limit neural noise or artifactual activity, and are used to improve signals quality prior to the extraction of different brain features. One popular approach in the literature involves the decomposition of brain signals into distinct frequency bands (i.e., delta, theta, alpha, beta, gamma and high-gamma) broadly divided into low-frequency components (LFC, below 40Hz) and high-frequency activity. These different frequency bands have been extensively used as features to model brain phenomena [1, 2, 3, 4, 5, 6, 7, 8, 9, 10, 11, 12]. For instance, LFC usually measured as the raw brain signal low-pass filtered (below 40 Hz)[13, 14, 15], have been used in many applications, including decoding of position, velocity and acceleration of executed motor tasks [16, 17, 18, 19, 20, 21, 13, 14, 15] leading to improvements compared to models restricted to amplitude modulations in only the alpha and beta bands [22, 19, 20]. Given the enhanced signal to noise ratio compared with both EEG and MEG, ECoG can also exploit high-frequency components above 100 Hz. High frequency band activity (HFB), usually measured as the averaged power changes in the band from 70 to 200 Hz, has been used for decoding in multiple tasks, including motor, auditory, and visual [16, 17, 18, 19, 20]. The extracted features are used to model brain responses for basic brain research, medical diagnostic, and rehabilitation areas. In rehabilitation, brain signals are used to control external devices that allow subjects to interact with the environment. In this case, a successful use of the device requires modeling the relationship between the executed task/stimulus and the brain signal. Models based on multilinear regression, support vector machines, probabilistic graphical models, and artificial neural networks have been proposed in combination with different types of features [23, 24, 3, 4, 6, 7, 8, 9, 17, 18, 19, 20, 21, 13, 14, 15]. These features involve spatial patterns discovery [25, 26], temporal modeling through probabilistic approaches such as hidden Markov models [27, 28], conditional random fields [16, 6, 7, 8, 9] and recurrent neural networks [29]. In addition, features based on frequency decomposition of brain signals performed through either Fourier or wavelet transform are well described in the literature. Each methods presents advantages and disadvantages. In multilinear regression, a preselection of the features is required to obtain good results, and incorporation of information about time lags between the brain response and the stimulus presented should be added manually introducing extra terms that are delayed versions of the actual brain response [30, 31, 18]. Without appropriate regularization this can introduce model over-fitting. In probabilistic graphical models and deep neural networks architectures, temporal relationships can be incorporated through modeling of long-range dependencies [16, 29]. However, these approaches require the use of a considerable amount of data that is usually not available in experiments with humans, limiting the performance of these methods [32].

In this work, we propose a method based on complex coherence to extract relevant information from the brain signals aiming to predict the presented stimulus (or dynamics of the executed task). The method is built on the idea that each experimental trial is a realization of a random process. The proposed approach takes into account different parameters such as frequency, phase and spatial distribution of brain signals, hence avoiding the need to predefine the frequency bands of interest or the phase relationships (i.e., lags) between stimulus and brain responses.

## 2. Methods

### 2.1. General description of the proposed method

The proposed method is described in Figure 1 using a finger movement task as an example. To model brain responses to stimulus/task-execution, each trial of an experiment is assumed to be a realization of a random process. Individual trials measure the combination of the response due to the stimulus/task plus random noise that is assumed to be uncorrelated to the actual brain response of interest. The proposed method makes use of complex coherence to determine the relationship in amplitude and phase between stimuli and brain signals at each frequency components across the training trials. In this approach the coherence is estimated across trials, instead of across time windows in each trial. This allows relaxing assumptions about the stationarity of the brain signals and is performed independently for each electrode, producing a spectral filter per recording site.

**Figure 1:**
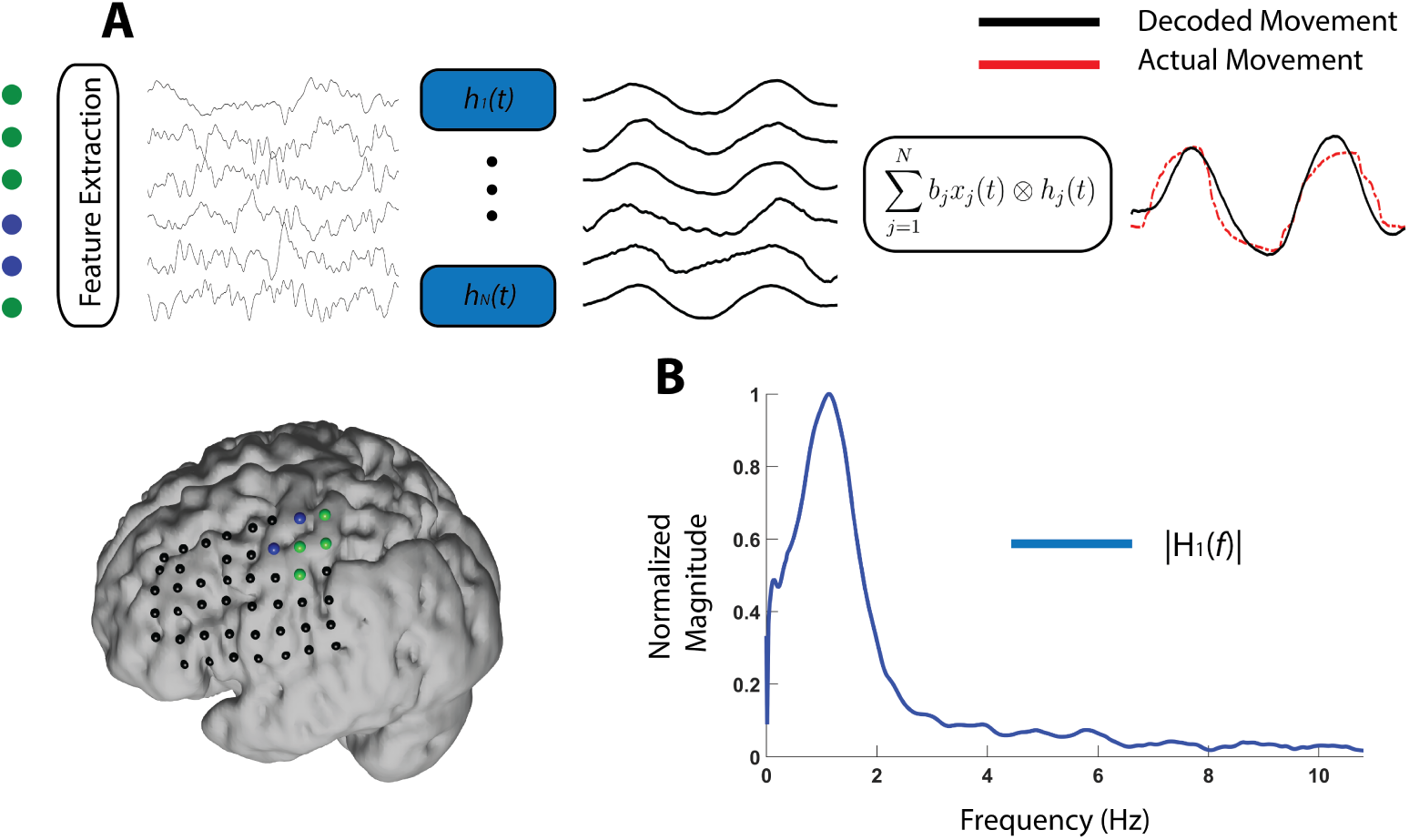
Description of the coherence-based spectro-spatial filter. The diagram represents the recordings of a trial while the subject perform a movement with his thumb. **A.** Brain signals recorded from electrodes in M1 (blue) and S1 (green) are used as input to a set of linear filters trained for each electrode. The filters represent the transfer function of a linear system that maps the brain signals to the movement of the finger. The outputs of all filters are combined using linear regression to produce the final prediction of the movement. The signals shown in the figure correspond to LFCs components. **B.** Example of the magnitude response of one of the learned filters showing a pick in the low-frequency domain around 1Hz.

The spectral filter extracts commonalities between stimuli and brain signals at each frequency band. The output of these filters is then combined using multilinear regression, producing a final prediction that incorporates spectral and spatial features of the brain response. Importantly, spatial filtering results can be analyzed to determine the contribution of different recording sites on the prediction, examining for anatomical discriminability among different cognitive tasks or different stimuli.

### 2.2. Finger movement data-set description

The motor data-set consists of electrocorticographic recordings from nine patients executing a repetitive finger movement task. During the task, subjects were cued with a word displayed on a bedside monitor indicating which finger to move (Thumb, Index, Middle, Ring, and Pinky). Subjects were asked to repetitively move the indicated finger during an interval of 2-second (trial). There were thirty trials for each finger. In addition to ECoG, finger positions were recorded using a 5-degree-of-freedom data-glove sensor. The data-glove signals were originally sampled at 25Hz and up-sampled at 1000Hz to match the sampling rate of the ECoG signals. The data-glove signal for each finger is the signal to be predicted from the brain signals. Two subjects without electrode coverage in the sensory-motor (S1) and Motor areas (M1) were excluded.

### 2.3. Speech perception data-set description

The data-set consists of ECoG recordings from three subjects. During the experiment, subjects we requested to listen to a taped female voice repeating six different words. Each trial started with a baseline period of 500ms after which a word out of a total of six is randomly selected and played on speakers at the bedside of the subject. Based on anatomical mapping, electrodes that responded to auditory stimulus were selected. Each word was repeated eighteen times. The audio input was recorded in parallel with brain signals to achieve minimum loss of synchronization, and all signals were sampled at 9600 Hz to cover the important portions of the voice spectrum. We use the speech envelope [33, 34] as the feature to be predicted from brain signals using the proposed method.

### 2.4. Speech production data-set description

The data-set consists of ECoG recordings from three subjects (same subjects as in the perception task). During the experiment, subjects repeated a particular word presented to them (among 6 different words). Each trial started with a baseline period of 500 ms after which the subject repeated the word that he or she heard prior to the beginning of the trial. Based on anatomical mapping, electrodes that responded to auditory stimulus and speech production were selected. Each word was repeated eighteen times. Technical details of the recordings are the same as described in the speech perception data-set. Because the subject decided when to speak, the data contains high variability regarding the onset of the speech production.

### 2.5. Ethics Statement

Finger movement data-set: All patients participated in a purely voluntary manner, after providing informed written consent, under experimental protocols approved by the Institutional Review Board of the University of Washington (#12193). All patient data was anonymized according to IRB protocol, in accordance with HIPAA mandate. These data originally appeared in the manuscript “Human Motor Cortical Activity Is Selectively Phase-Entrained on Underlying Rhythms” published in PLoS Computational Biology in 2012 [35].

Speech perception and speech production data-set: All patients volunteered and gave their informed consent (experimental protocol was approved by the Albany Medical College Institutional Review Board and methods were carried out in accordance with the approved guidelines and regulations) before testing. These data originally appeared in [36].

### 2.6. Signal Preprocessing and Feature Extraction

For all three data-sets the preprocessing stage is identical. All the electrodes were band-pass filtered between 0.1 and 200 Hz and a notch filter at 60 Hz with additional filtering of harmonics at 120 and 180 Hz reduced the interference of the power line. A common average reference (CAR) was used to reduce the effect of the reference electrode in all the recording sites. For this, the average across all electrodes is subtracted from the individual electrodes in the following fashion:

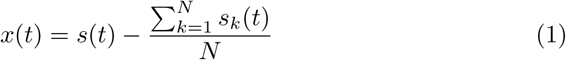

where *s*(*t*) = *{s*1(*t*), …, *s*_*N*_ (*t*)} represent the raw signals measured at different electrode locations, and *N* is the number of electrodes.

The brain signals were separated within two frequency ranges: low frequency components (LFC) from 0.5 to 40Hz, and HFB(70 to 170 Hz). LFCs were estimated by filtering the brain signals with a band-pass Butterworth filter of 4th order between 0.5 and 40 Hz. The envelope of high frequency band signals (HFBE) was calculated as the magnitude of the analytic signal in the following fashion: assuming that the signal *x*_*bandpass*_(*t*) is the result of band-pass filtering *x*(*t*) in the HFB range, the analytic signal *x*_*analytic*_(*t*) is calculated as:

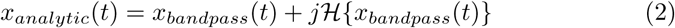

where *ℋ* represents the Hilbert transform operation. The envelope of high gamma was low-pass filtered with a Butterworth filter of 4^*th*^ order and a cut frequency of 40 Hz to reduce rapid changes in the amplitude of the signal.

### 2.7. Coherence-based Spectro-Spatial Filter

The complex coherence allows for determining how well two signals correlate at each frequency component [37]. Given the random variables *x*(*t*) = {*x*1(*t*), …, *x*_*N*_ (*t*)} representing the brain features extracted at each recording electrode and *y*(*t*) representing the dynamics of the stimuli that elicits the brain responses, the complex coherence between each *x*_*j*_(*t*) and *y*(*t*) is given by:

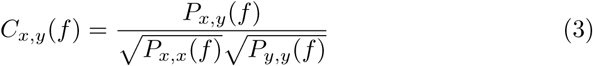

where *P*_*x,x*_(*f*) and *P*_*y,y*_(*f*) are the power spectral density of *x*_*j*_(*t*) (brain signal at electrode *j*) and *y*(*t*) respectively, and *P*_*x,y*_(*f*) is the cross-power spectral density calculated between *x*_*j*_(*t*) and *y*(*t*). The magnitude squared of the complex coherence has values between 0 and 1 and can be understood as the squared correlation between the two signals at each frequency component. Using the coherence as a measure of correlation between two signals at each frequency *f*, a linear filter is defined as:

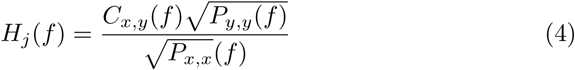

Using the expression in Equation 3 for coherence we obtain:

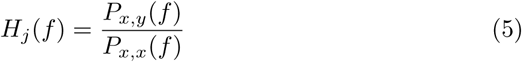

where *H*_*j*_(*f*) is the linear filter. Note that *P*_*x,y*_(*f*) is a complex quantity, while *P*_*x,x*_ is real. Therefore the phase spectrum of *H*_*j*_(*f*) is given by the phase differences between *y*(*t*) and *x*_*j*_(*t*). That is, for prediction of *y*(*t*) from *x_j_*(*t*), the phase differences at each frequency *f* are taken into consideration by the filter. Estimation of *y*(*t*) from *x_j_*(*t*) is then obtained by:

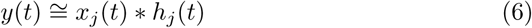

where *h*_*j*_(*t*) represent the inverse Fourier transform of *H*_*j*_(*f*). Finally, different electrodes may contain different type of information, and should be accordingly combined forming an spatial filter, as follows:

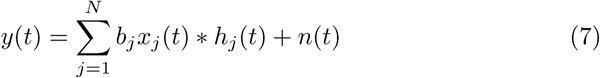

where *b_j_* is a weight that determines how important is the signal coming from the *j*^*th*^ electrode for the prediction of the signal *y*(*t*). The term *n*(*t*) is used to model the error in the prediction. The set of coefficients *b_j_*, can be understood as a spatial filter that provides information about which brain areas are involved in the processing of the stimuli or task executed. Combining *h*(*t*) with the coefficients in Equation 7 forms a filter that takes into account the frequency and phase spectrum of the signals, and the spatial patterns representing the contribution of different brain areas.

Filters *h*_*j*_(*t*) are build by calculating the coherence and power spectral densities using the Welch Method [38] assuming each trial as a realization of a random process. It is worth noting that the frequency response of the filters *h*(*t*) is defined by the signals *x*_*j*_(*t*) and *y*(*t*), and therefore should be calculated independently for each subject. Once the filters *h_j_*(*t*) are build, parameters *b*_*j*_ can be learned using the least square solution for linear regression. For validation of the performance of the method, the proposed filters are constructed using portion of the available data (training set) and tested in the remaining portion (testing set), which is never seen during the training stage.

### 2.8. Methods used for comparison

We implemented two methods, which we compared against the coherence-based spectro-spatial filter. Multilinear regression model was fitted to the data using LFCs and HFBE features. As the linear regression method cannot account for the latency between brain signals and tasks/stimuli, the brain signal features were lagged using lag values in a windows of 0 to 500 ms.

A second method used for comparison is a non-linear fitting strategy based on artificial neural networks (named ANN-fit hereafter). ANN-fit model contains a hidden layer of one hundred neurons. Neurons in the hidden layer have a sigmoid activation function while the output layer uses a linear function. As for the multilinear method, the feature set was expanded including lagged versions of the original features. Both methods, multilinear regression and ANN-fit, degrade without the incorporation of lagged regressors. Importantly, the proposed coherence-based spectro-spatial filters method does not require the incorporation of any lags for the features as this is computed automatically during the training stage by measuring phase differences between the brain signals and the stimulus to predict.

## 3. Results

To address whether the coherence-based spectro-spatial filters method out-performs traditional approaches, we first compared its predictive power (in terms of correlation between the predicted output and the actual stimulus/task dynamics) to a multilinear regression model as well as to a model based on artificial neural networks as described in Section 2.8. Evaluation of all methods was performed in the same fashion, with the same features, using leave-one-out cross-validation. Figure 2 shows the results averaged across all subjects for the three data-sets used.

**Figure 2:**
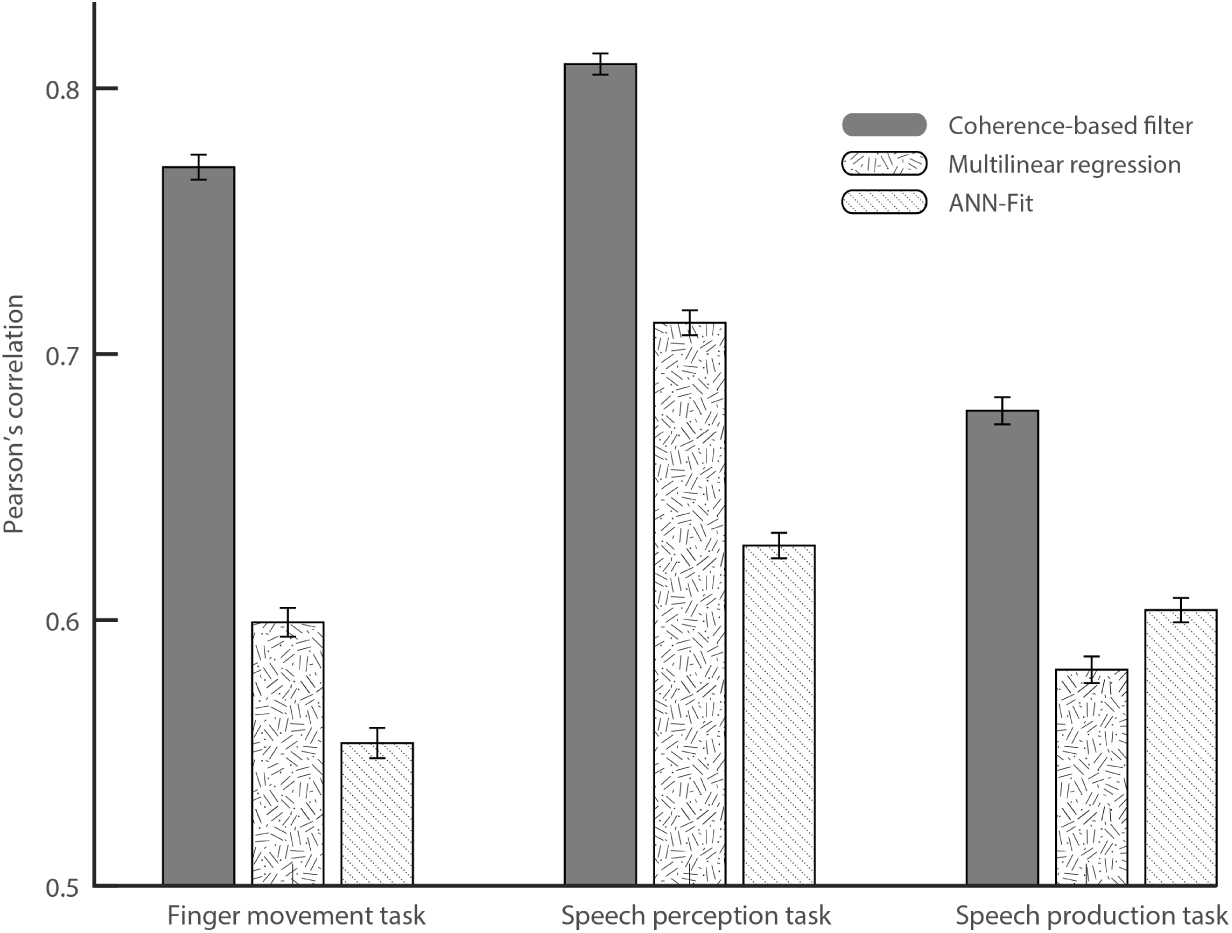
Average Pearson’s correlation for three different tasks using three different methods. Error bars correspond to standard error of the mean.

To assess statistical significance of performance gain, we fitted a generalized linear model (GLM) using as response variable the correlation obtained between the actual and the predicted output for each trial. The effects are the types of model (3 levels), the conditions (5 levels for the motor movement data-set, and 6 levels for speech perception and production data-sets) and the subjects (7 levels for the motor movement data-set and 3 levels for the other data-sets). The contrast analysis of the coefficients of the GLM (the type of model effects) shows that the proposed method provides significant improvement (*p <* 0.05) over both the multilinear model and the ANN-fit model for the finger movements, the speech perception and the speech production modalities. In addition, no significant differences were found between the multilinear regression and the ANN-fit model at 95% confidence level. Importantly, the improved correlation values between the actual and predicted feature dynamics for the proposed method was observed in all subjects across the three modalities (see Tables 1, Table 2 and 3 for details).

**Table 1:** Averaged correlation values between the predicted and the real finger movement dynamics for the three models.

| Subject | Proposed Method |  |  | Linear Regression | ANN-fit |
| --- | --- | --- | --- | --- | --- |
|  | LFC | HFBE | LFC+HFBE | LFC+HFBE | LFC+HFBE |
| S01 | 0.80 | 0.70 | **0.80** | 0.71 | 0.67 |
| S02 | 0.81 | 0.74 | **0.82** | 0.68 | 0.63 |
| S03 | 0.79 | 0.54 | **0.81** | 0.58 | 0.53 |
| S04 | 0.76 | 0.56 | **0.80** | 0.55 | 0.52 |
| S05 | 0.72 | 0.56 | **0.75** | 0.64 | 0.62 |
| S06 | 0.62 | 0.43 | **0.63** | 0.44 | 0.39 |
| S07 | 0.71 | 0.68 | **0.78** | 0.57 | 0.51 |
| Average | 0.74 | 0.60 | **0.77** | 0.60 | 0.55 |

**Table 2:** Averaged correlation values between the predicted and the real speech envelope for the three models.

| Subject | Proposed Method |  |  | Linear Regression | ANN-fit |
| --- | --- | --- | --- | --- | --- |
|  | LFC | HFBE | LFC+HFBE | LFC+HFBE | LFC+HFBE |
| S01 | 0.82 | 0.77 | **0.83** | 0.71 | 0.62 |
| S02 | 0.75 | 0.68 | **0.74** | 0.64 | 0.55 |
| S03 | 0.83 | 0.75 | **0.86** | 0.78 | 0.70 |
| Average | 0.80 | 0.73 | **0.81** | 0.71 | 0.62 |

**Table 3:** Averaged correlation values between the predicted and the real of the envelope of speech production for the three models.

| Subject | Proposed Method |  |  | Linear Regression | ANN-fit |
| --- | --- | --- | --- | --- | --- |
|  | LFC | HFBE | LFC+HFBE | LFC+HFBE | LFC+HFBE |
| S01 | 0.56 | 0.70 | **0.67** | 0.56 | 0.54 |
| S02 | 0.72 | 0.61 | **0.66** | 0.60 | 0.63 |
| S03 | 0.69 | 0.72 | **0.72** | 0.58 | 0.64 |
| Average | 0.66 | 0.67 | **0.68** | 0.58 | 0.60 |

Furthermore, the proposed method enables to analyze the importance or weight of different recording sites in the prediction of the stimulus/task dynamics. As a result, the magnitude of coefficients *b_j_* (see Equation 7) reflects the contribution of each electrode to the final prediction. In the proposed method, the combination of channels occurs after the filtering stage. Therefore, the output of each filter only contains components that are linearly related to the stimulus/task dynamics. Consequently, the set of coefficients *b_j_* can be understood as a set of spatial filters that contain discriminant information about the areas involved in task execution. These spatial filters can be plotted on the brain models for different tasks. Figure 3 displays the spatial distribution of the coefficients for LFC (panel B) and HFBE (panel A) components in one representative subject of the finger movement data-set. The results show that the proposed method leads to distinct spatial patterns, not only in response to different finger movements but also in the LFC and HFBE components. Similarly, the spatial filter for predicting the envelope of the perceived (Figure 4) and produced (Figure 5) speech revealed a level of discriminability across words as well as between the low and high-frequency components.

**Figure 3:**
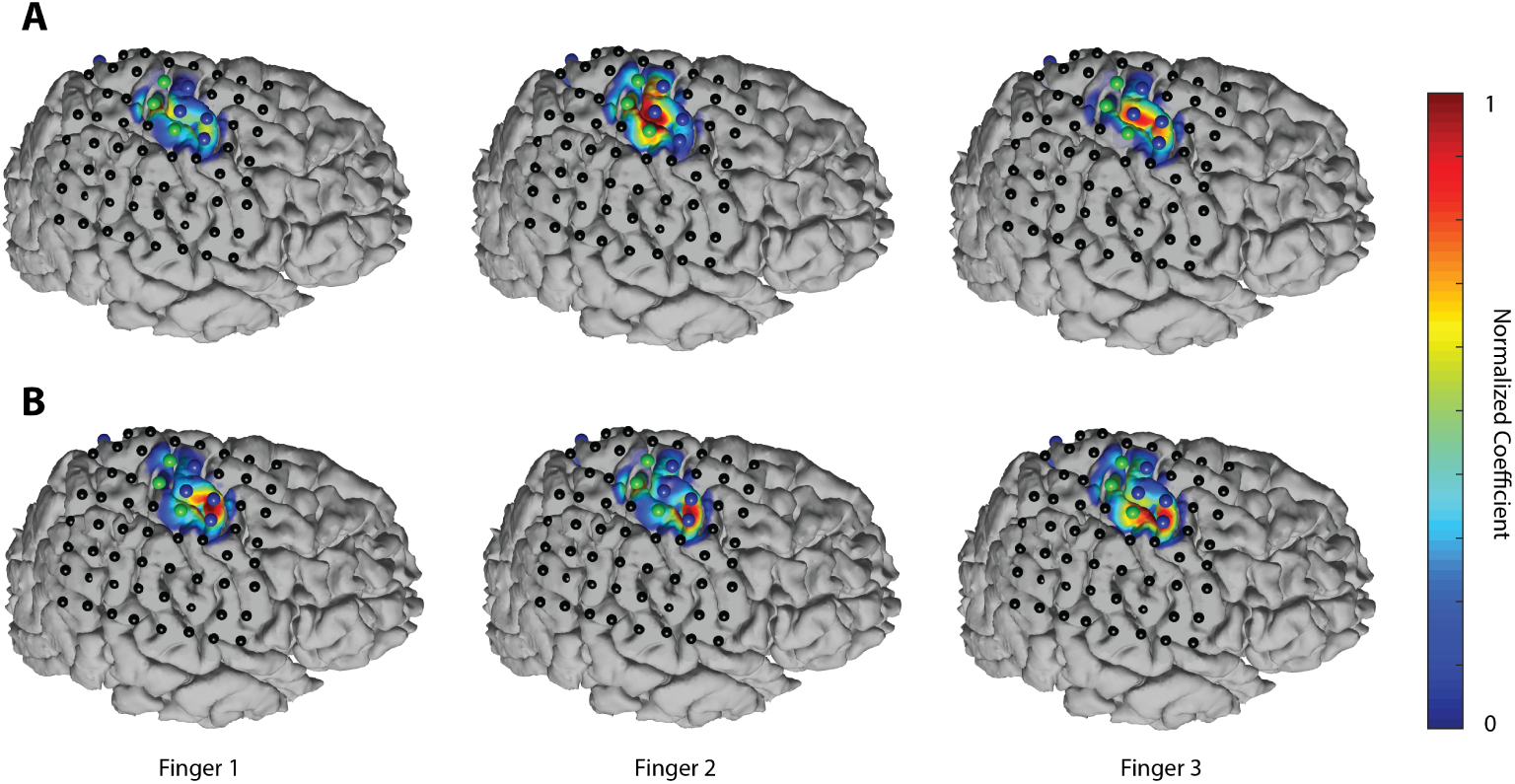
**A.** Spatial patterns of (A) LFC and (B) HFBE of a representative subject during movement of three different fingers.

**Figure 4:**
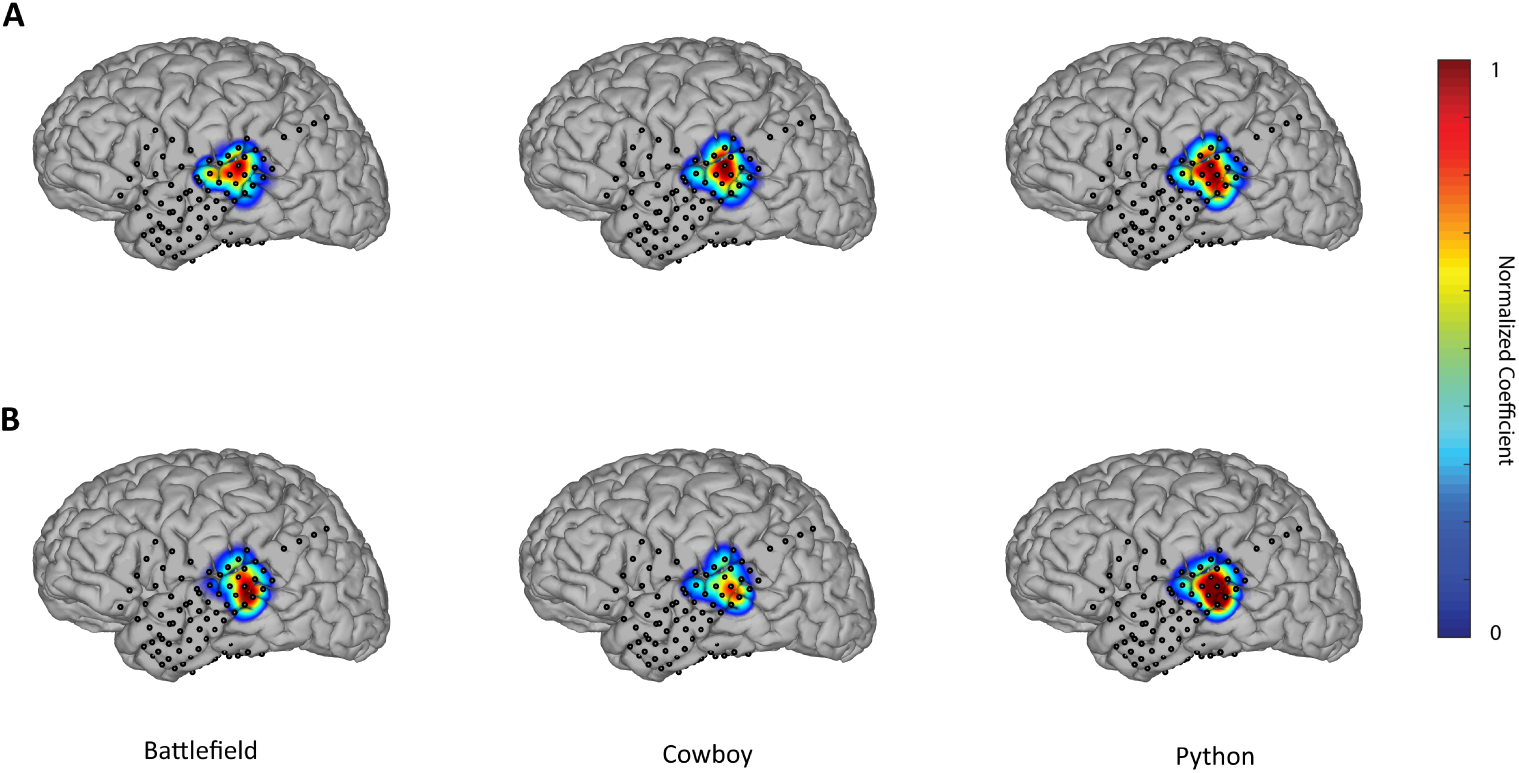
**A.** Spatial patterns of (A) LFC and (B) HFBE of a representative subject during the perception of three different words.

**Figure 5:**
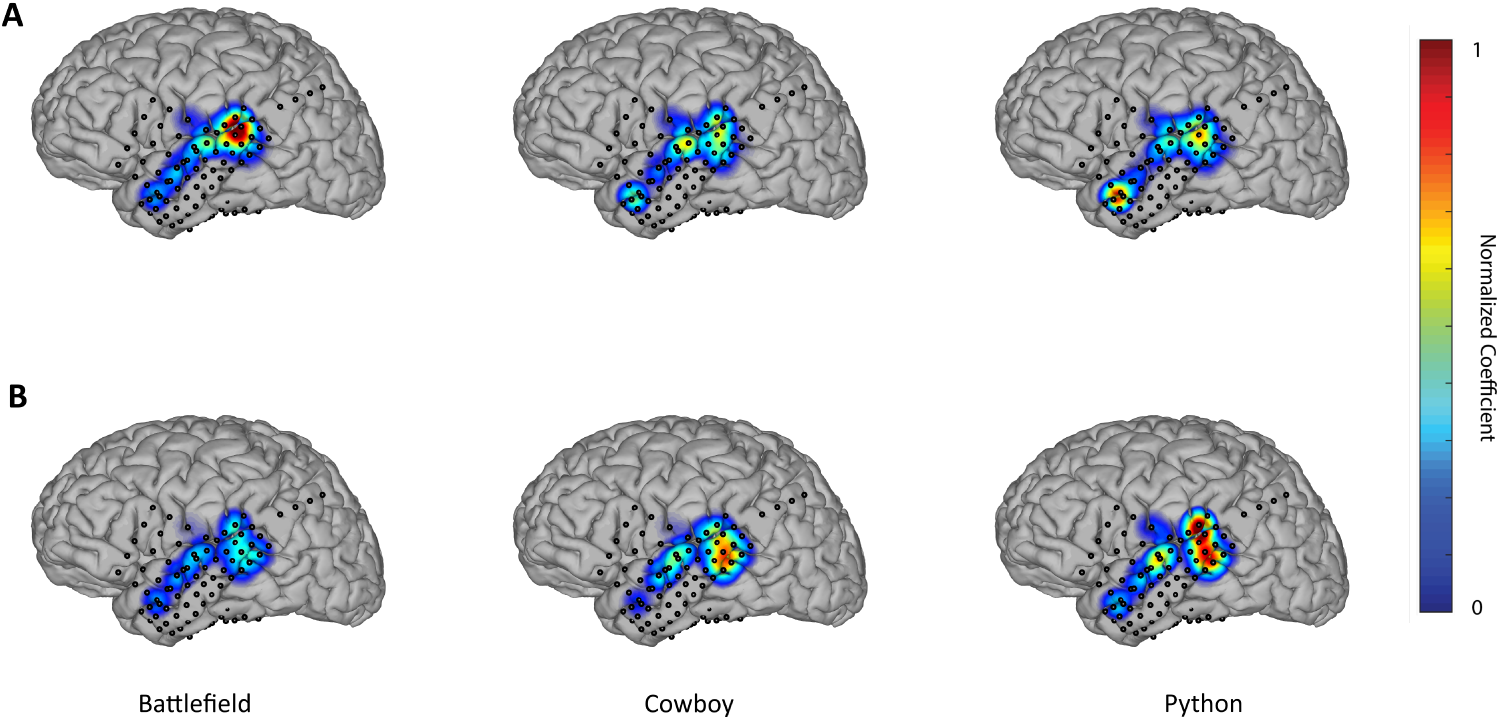
**A.** Spatial patterns of (A) LFC and (B) HFBE of a representative subject during the production of three different words.

To assess the degree of discriminability within conditions (finger moved or word perceived/produced) revealed by the magnitude of the weights *b*_*j*_, we trained a set of models for each condition using a leave-one-out cross-validation approach. To address this, we ranked the level of discriminability of the co-efficients *b*_*j*_ assigned to each electrode using K-means and fed an LDA (linear discriminant analysis) classifier with only the best two electrodes (selected using only the training set) to prevent overfitting. The results revealed that the coefficients of the best two electrodes enabled discrimination of the models fitted to each finger movements with a high-level of accuracy for both LFCs (83%, std = 0.16) and HFB (80%, std = 0.06). Similarly, we found that the coefficients discriminated among the models learned for different words in the speech perception (LFCs 90%, std = 0.07; HFBE 94%, std = 0.04) and production (LFCs 88%, std = 0.11; HFBE 93%, std = 0.05) tasks. These results show that the method learns for each condition a spatial filter that indicates the importance of a particular electrode in the decoding of the task dynamics.

## Discussion

### 3.1. Coherence-based spectro-spatial filter

We propose and assess a method to reconstruct stimuli or task dynamics from brain recordings that does not require a priori specification of any signal parameters. We found that the coherence-based method outperforms traditional predictive models and provides both a high-performance level (in terms of correlation), and consistency across motor and linguistic domains. Notably, we demonstrate that the coherence-based method produces spatial filters that are discriminative of the task executed, revealing the importance of different brain areas for the execution of the tasks. The performance improvement relative to other methods is explained by several factors. The proposed method uses frequency decomposition and takes into account the phase relationships across trials. This allows for removing frequency components containing artifactual high power that do not show phase consistency between stimulus and brain signals across trials. Such phase relationships are an essential factor because they reflect the latencies between the signals of interest at each frequency component. Multilinear regression cannot account for phase relationships and requires expanding the set of regressors with lagged versions of the data. Although the ANN-fit method includes the possibility to incorporate nonlinearities through sigmoid activation functions, it cannot model latencies among the signals of interest. And even though expanding the regressors set with lagged versions of the data is possible as presented here and in [24, 30, 31, 18], this may lead to over-fitting as the number of parameters increases while the amount of available data remains constant.

The proposed coherence-based filter shows robust performances across different subjects and operates well regardless of interindividual differences and electrodes localization making it well suited for cases where the decoding goal is to decode what stimulus was presented to the subject. Using complex coherence, the correlations between stimulus and brain signals at each frequency component are learned and are used to create a filter. The filter parameters are calculated under the assumption that each trial is a realization of a random process which allows employing the different trials to calculate a robust estimation of the cross-spectral density between brain signals and the stimulus presented. This permits extraction of components that are in phase with task dynamics across trials for each frequency and recording site. The resulting signal is then combined spatially to form a final prediction. The coherence-based spectro-spatial filter method has the advantage of including different dimensions of brain signals such as phase, frequency and space to handle prediction of stimulus dynamics in an automatic fashion. Importantly, the second stage of the proposed method combines different recordings sites. The difference between this and the methods used for comparison is that signals are combined after the filtering, which ensures that the components combined have a linear relationship with the stimulus/task dynamics. This reduces the possibility that the coefficients learned (spatial filters) reflect a simple noise canceling process. Evidence presented in this study on clustering of the values of the coefficients for each task enables separating the models learned by stimulus type (word presented) or task (finger moved) providing information about the brain areas involved in the particular task.

### 3.2. Caveats and caution

The proposed method is well suited for discrete tasks. For instance, speech perception experiments as those presented here, in which the subject listens to a word and a prediction of the acoustic envelope of the audio attended is made, is an excellent example of such a discrete task. The proposed method could be used for continuous prediction as long as the causality of the filters *h*_*j*_(*t*) is ensured. Although engineering methods for ensuring causality exists, particular implementations of these techniques are beyond the scope of this study. Finally, the spatial filters could lead to over-fitting if too many electrodes are used, or if the data-set is too small. This issue was handled in this study by selecting areas according to prior information from anatomical mapping performed on the subjects. However, this can be controlled automatically including regularization parameters that introduce sparsity on the model effectively minimizing the risk of over-fitting if the amount of data available allows for it.

## 4. Conclusion

We present a method capable of predicting from brain signals, characteristic features of a stimulus. The proposed method employs complex coherence to extract common patterns among the brain signals related to the dynamics of the presented stimulus. This includes spatial information forming a spectro-spatial filter that is capable of reconstructing the dynamics of the stimulus with high performance (in terms of correlation coefficient). Analysis of the coefficients that form the learned spatial patterns showed discriminability among different conditions, indicating the involvements of different areas and frequency components during the execution of various cognitive tasks such as finger movement as well as speech perception and production. The anatomical discriminability revealed by the method can be exploited in the design of neuro-prosthesis as well as for investigating the normal brain function.

